# Premature arterial stiffening in Hutchinson-Gilford Progeria Syndrome linked to early induction of Lysyl Oxidase (LOX) and corrected by LOX inhibition

**DOI:** 10.1101/773184

**Authors:** Ryan von Kleeck, Sonja A. Brankovic, Ian Roberts, Elizabeth A. Hawthorne, Kyle Bruun, Paola Castagnino, Richard K. Assoian

**Affiliations:** Department of Systems Pharmacology and Translational Therapeutics, University of Pennsylvania, Philadelphia PA 19104; Institute of Translational Medicine and Therapeutics, University of Pennsylvania, Philadelphia PA 19104; Center for Engineering MechanoBiology, University of Pennsylvania, Philadelphia PA 19104; G.W. Woodruff School of Mechanical Engineering, Georgia Tech, Atlanta GA 30313

## Abstract

Arterial stiffening is a hallmark of premature aging in Hutchinson-Gilford Progeria Syndrome (HGPS), but the key molecular regulators initiating arterial stiffening in HGPS remain unknown. To identify these early events, we compared arterial mechanics and ECM remodeling in very young HGPS (LMNA^G609G/G609G^) mice to those of age-matched and much older wild-type (WT) mice. Biaxial inflation-extension tests of carotid arteries of 2-month mice showed that circumferential stiffness of HGPS arteries was comparable to that of 24-month WT controls whereas axial arterial stiffening, an additional hallmark of normal aging, was mostly spared in HGPS. In an effort to identify underlying mechanisms, we examined expression levels of the major stiffness-regulatory molecules in WT and HGPS arteries. Transmission electron microscopy revealed slightly increased amounts of collagen within the elastin folds of HGPS carotid arteries, but this change was barely detectable by immunostaining carotid cross sections or qPCR of isolated aortas for collagens I, III, or V. Elastin integrity was also similar in the WT and HGPS arteries. In contrast, immunostaining readily revealed an increased expression of Lysyl oxidase (LOX) protein in young HGPS carotid arteries relative to aged-matched WT controls. Further analysis showed that HGPS arteries express increased amounts of LOX mRNA, and this effect extends to each of the arterial LOX family members. Remarkably, treatment of HGPS mice with the pan-LOX inhibitor β-aminopropionitrile (BAPN) restored near-normal circumferential arterial mechanics to HGPS carotid arteries, mechanistically and causally linking LOX upregulation to premature arterial stiffening in HGPS. Finally, we show that this premature increase in arterial LOX expression in HGPS foreshadows the increased expression of LOX that accompanies circumferential arterial stiffening during normal aging.

## INTRODUCTION

Hutchinson-Gilford Progeria Syndrome (HGPS) is a rare genetic disease of premature aging. HGPS is caused by an autosomal dominant mutation in *LMNA*, the gene encoding Lamin A (Gonzalo et al. 2016; Capell et al. 2007). The mutation is “silent” (LaminA^G608G^) but leads to a defect in biosynthetic processing of the Lamin A precursor and results in a truncated, farnesylated protein called progerin, that is defective in its localization within the nucleus (Gonzalo et al. 2016; Capell et al. 2007). Children with HGPS typically die in their teenage years as a consequence of cardiovascular disease (atherosclerosis, myocardial infarction and/or stroke) (Gonzalo et al. 2016; Capell et al. 2007). These cardiovascular consequences occur in the absence of high cholesterol or triglycerides (Gordon et al. 2005), suggesting that cholesterol-independent risk factors must be at play. Intriguingly, the arteries of HGPS patients are abnormally stiff as indicated by increased pulse-wave velocity (Gerhard-Herman et al. 2011), and arterial stiffness is a cholesterol-independent risk factor for cardiovascular disease (Mitchell et al. 2010). Pathologically stiff arteries lead to increased load on the heart (Safar 2010; Laurent & Boutouyrie 2015), which can then have systemic consequences. Despite its importance, the mechanical properties of HGPS arteries and how progerin expression leads to increased arterial stiffness remain poorly understood. Interestingly, the vascular pathology of HGPS has several similarities to the vascular pathology of naturally aged arteries (Olive et al. 2010). Indeed, increased arterial stiffness is a hallmark of normal aging and increases substantially after the age of 50 in healthy males and females (Mitchell et al. 2007). However, the arterial stiffness of HGPS children resembles that of ~60 year-old individuals (Gerhard-Herman et al. 2011), indicating a striking acceleration of arterial stiffening in HGPS.

The composition of the extracellular matrix (ECM) plays a critical role in arterial stiffness (Cox & Erler 2011). The stiffness of the major arteries relies on a balance of elastin and collagen fibers (Humphrey et al. 2014) to maintain a proper response to blood pressure. Elastin allows for recoil at lower load and dampens cyclic loading from the beating heart while fibrillar collagens contribute to the strain-stiffening property of arteries at high load (Kohn et al. 2015). Arteries express three main fibrillar collagens, with collagen-I>collagen-III>collagen-V in abundance (Hulmes 2008; Prockop & Kivirikko 1995). Increased collagen deposition as well as elastin fragmentation has been observed in the arteries of HGPS children at autopsy (Olive et al. 2010; Gerhard-Herman et al. 2011; Merideth et al. 2008) and in aged arteries (Kohn et al. 2015; Tsamis et al. 2013).

In addition to increases in fibrillar collagens, the mechanical properties of tissues are further regulated by matrix remodeling enzymes (Freitas-Rodríguez et al. 2017; Lampi & Reinhart-King 2018), and especially Lysyl Oxidase (LOX) and its family members (LOXL1, LOXL2, LOXL3 and LOXL4) (Yamauchi & Sricholpech 2012; Nilsson et al. 2016; Steppan et al. 2019). These enzymes covalently cross-link newly synthesized collagen and elastin fibers to enhance their tensile strength (Tsamis et al. 2013; Baker et al. 2013; Herchenhan et al. 2015), and their overexpression is commonly seen in stiffness-related pathologies (Desmoulière et al. 1997; Kagan 1994). However, since elastin biosynthesis ends early in life (Tsamis et al. 2013; Wagenseil & Mecham 2012; Davidson et al. 1986; Mithieux & Weiss 2005; Kelleher et al. 2004), LOX-mediated crosslinking is thought to target newly synthesized collagens in normal aging.

HGPS is a very rare genetic disease (Gerhard-Herman et al. 2011; Olive et al. 2010) and since the number of HGPS patients is so small, the detailed analysis of early and potentially causal events in the pathogenesis of HGPS relies heavily on animal models. Osorio et al. (Osorio et al. 2011) developed a Lmna^G609G/G609G^ knock-in mouse (hereafter called HGPS mice) that models the mutation in human HGPS and displays many traits of the human disease including premature death. An increase in the expression of several collagens, including collagen-III, IV, and V is also apparent in these mice as they reach 3-4 months (del Campo et al. 2019), an age that is toward the end of the HGPS mouse lifespan (Osorio et al. 2011). However, whether these changes in arterial stiffness are primary or secondary effects of HGPS is not known.

In this report, we have studied very young HGPS mice as well as age-matched and much older WT mice to characterize the early arterial stiffening that occurs well before morbidity and death in HGPS. Using biaxial inflation-extension tests with freshly isolated carotid arteries, we report that arterial stiffening in HGPS is strikingly premature but preferentially in the circumferential direction. To identify the corresponding mechanism(s), we examined the expression of mechanoregulatory ECM proteins in HGPS, tested their causal relationship to premature circumferential arterial stiffening, and determined if the same events might contribute to arterial stiffening in normal aging.

## RESULTS

### Premature arterial stiffening in HGPS mice

Our studies compared mechanics and ECM remodeling in the carotid arteries of WT and HGPS mice, as the carotid is a prominent site of atherosclerotic lesion formation, and its occlusion is thought to be responsible for induction of stroke in HGPS children (Gerhard-Herman et al. 2011). Freshly isolated arteries were stretched below, at, and above their *in vivo* lengths at constant pressure to provide insight into axial arterial stress (Fig. 1A). The artery was also stretched to its approximate *in vivo* length (referred to as the In Vivo Stretch; IVS), and the increase in outer diameter in response to incremental increases in pressure provided insight into circumferential vessel mechanics (Fig. 1D). These primary data were used to determine axial and circumferential stress-stretch relationships in which axial and circumferential stiffness is defined by the stress the artery experiences for a defined increase in stretch.

**Fig. 1.**
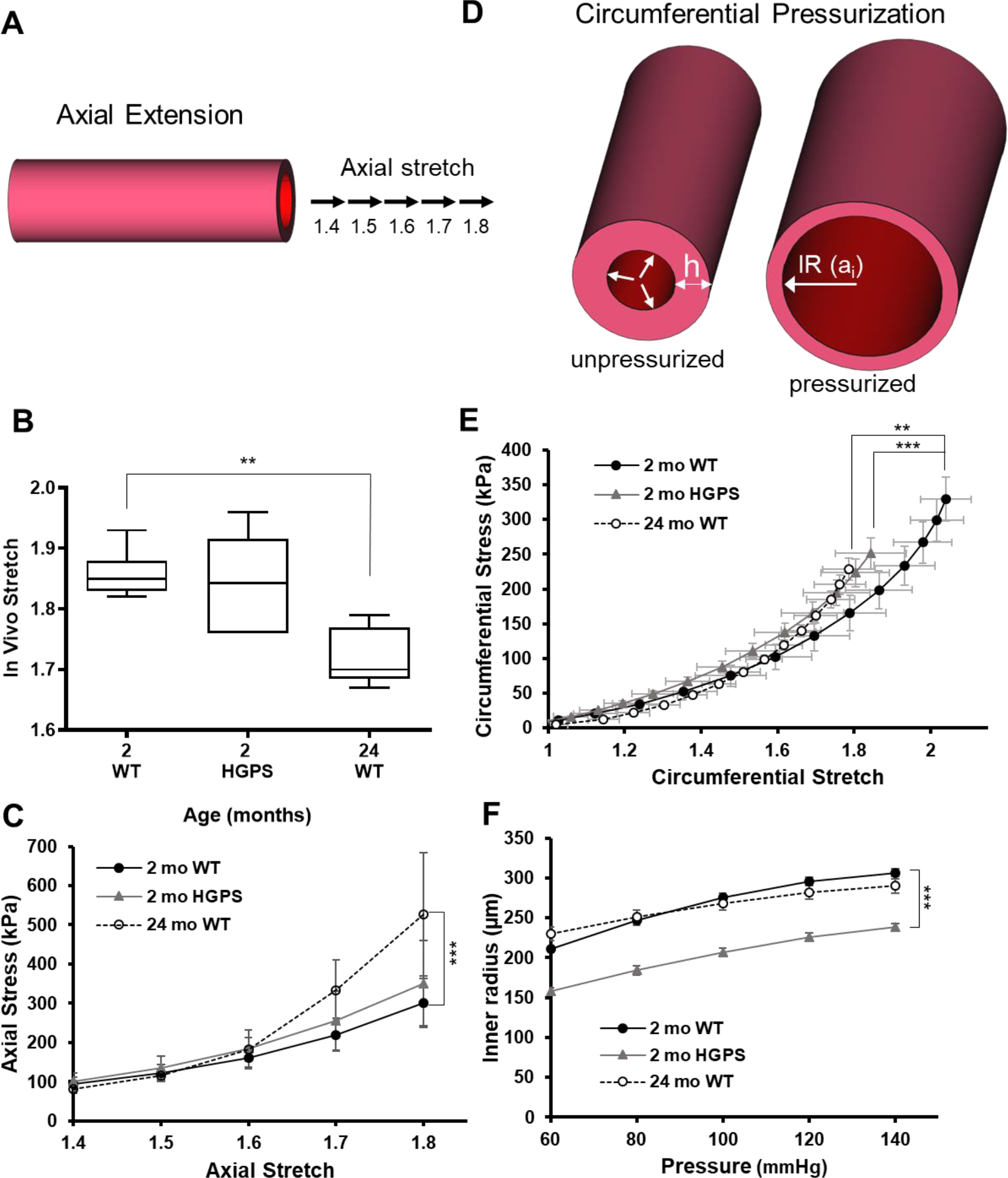
Mechanical properties of isolated carotid arteries display premature circumferential stiffening in HGPS mice. Carotid arteries from male 2-month (n=7) and 24-month (n=5) WT and 2-month HGPS (n=6) mice were analyzed by pressure myography. **(A)** Diagrammatic representation of the axial stretching protocol used to determine axial mechanics and “*In vivo stretch*.” **(B)***In vivo* stretch of 2-month and 24-month WT and 2-month HGPS arteries. Statistical significance was determined by Mann-Whitney relative to 2-month WT arteries. **(C)** Axial stress-stretch curve determined at 90-mm Hg for carotid arteries from 2- and 24-month WT mice and 2-month HGPS mice. Results show means ± SD. **(D)** Diagrammatic representation of circumferential arterial deformation with pressure. With pressurization, wall thickness (h) decreases while the loaded inner radius (a_i_) increases. **(E)** Circumferential stress-stretch curves of carotid arteries from 2- and 24-month WT mice and 2-month HGPS mice. **(F)** Changes in inner radius of 2- and 24-month WT and 2-month HGPS carotid arteries with pressure were determined as in Methods. Results in E and F show means ± SE. Statistical significance in panels C, E and F were determined by two-way ANOVA.

We recently used these and related mechanical analyses to examine the stiffening of carotid arteries of C57BL/6 mice aged from 2-24 months; the results showed decreases in IVS and increases in wall thickness as well as axial and circumferential arterial stiffness with age, beginning at 12-months and becoming even more evident by 24 months (Brankovic et al. 2019).Our studies here therefore compared arterial geometry and stiffness of young (2-month) WT and HGPS mice to those of 24-month WT mice to understand how arterial mechanics are affected in HGPS and how they compare to the changes that occur in normal aging.

Metrics of axial stiffening (IVS and axial stress-stretch curves) were surprisingly similar in 2-month WT and 2-month HGPS arteries (Fig. 1B and C, respectively). In contrast, the circumferential stress-stretch curve of 2-month HGPS carotid arteries, as derived from the changes in outer diameter with pressure (Fig. S1A), showed that circumferential arterial stiffness was similar to the stiffness of 24-month WT carotid arteries (Fig. 1E). Thus, arterial stiffening in HGPS mice is profoundly premature as it is in the human syndrome, but the early effect is anisotropic and selectively circumferential, an insight that has not been obtainable by pulse-wave velocity studies of HGPS children.

HGPS carotid arteries had significantly smaller inner radii than WT mice of either age (Fig, 1F). In contrast, carotid wall thickness of 2-month old HGPS mice was similar to 2-month WT controls and much less than the aged WT mice (Fig. S1B). This difference in vessel geometry between 2-month HGPS and 24-month WT carotids, despite similar circumferential stiffness, suggests that ECM remodeling in HGPS mice might be distinguishable from the arterial remodeling that accompanies normal aging.

The use of HGPS mice allowed for a detailed analysis of potential sex differences in arterial stiffness that has not been attainable given the small numbers of HGPS girls and boys. Like HGPS male mice, arterial compliance and circumferential stiffening was also premature in 2-month female HGPS mice (Fig. S2A-B), and the decrease in inner radius seen in males also held true for 2-month female HGPS mice relative to WT (Fig. S2C). As opposed to WT and HGPS males, the carotid arteries of 2-month female HGPS mice had a slightly reduced IVS and an increased axial stress as compared to 2-month WT females (compare Fig. S2D-E and Fig. 1). However, the difference in axial stress-stretch curves between WT and HGPS females reflected an overall reduction in axial stress of female mice as compared to males (compare Figs. 1B-C to S2D-E). HGPS females also show a statistically significant increases in wall thickness not seen in HGPS males (compare Fig. S1B and Fig. S2F).

Although reduced smooth muscle cell number has been reported in the ascending aorta of HGPS children at autopsy (Hamczyk & Andrés 2019), histological analysis of the 2-month WT and HGPS carotid arteries showed similar cellularity (Fig. S3A; H&E) and numbers of vascular smooth muscle cell (SMC) nuclei in the medial layer of young mouse carotids (Fig. S3B). Similarly, other markers of late HGPS vascular lesions including increases in calcium content (Merideth et al. 2008; Olive et al. 2010) and changes in elastin integrity were not seen in the HGPS carotid arteries at this early time-point (Figs. S3C and S4). We did not observe apoptotic cells in either WT or HGPS carotids (Fig. S3D) The absence of these late lesion markers supports the idea that our studies are examining initiating events in HGPS before the onset of overt HGPS phenotypes and that arterial stiffening is a very early event in disease progression. However, we did detect a small increase in the abundance of the senescence marker p16^INK4A^ in the medial layer of carotid arteries from 2-month HGPS mice as compared to the aged-matched WT controls (Fig. S3E-F).

### Canonical upregulation of fibrillar collagens is lacking in early HGPS

Tissue remodeling and stiffening typically involves increased amounts of fibrillar collagens and/or elastin fragmentation (see Introduction). Collagen-I is the major strain-stiffening component of the arterial ECM, but an immunostaining analysis of carotid sections was unable to detect statistically significant increases in collagen-I in either the medial or adventitial layers of 2-month HGPS carotid arteries as compared to age-matched WT controls (Fig. 2A-B and S5). Of the minor fibrillar collagens, these young HGPS carotid arteries showed a statistically significant but small increase in medial collagen-III and no change in the abundance of collagen-V (Fig. 2A-B and S5). Consistent with these results, none of the arterial fibrillar collagens showed increased mRNA expression in HGPS (Fig. 2C).

**Fig. 2.**
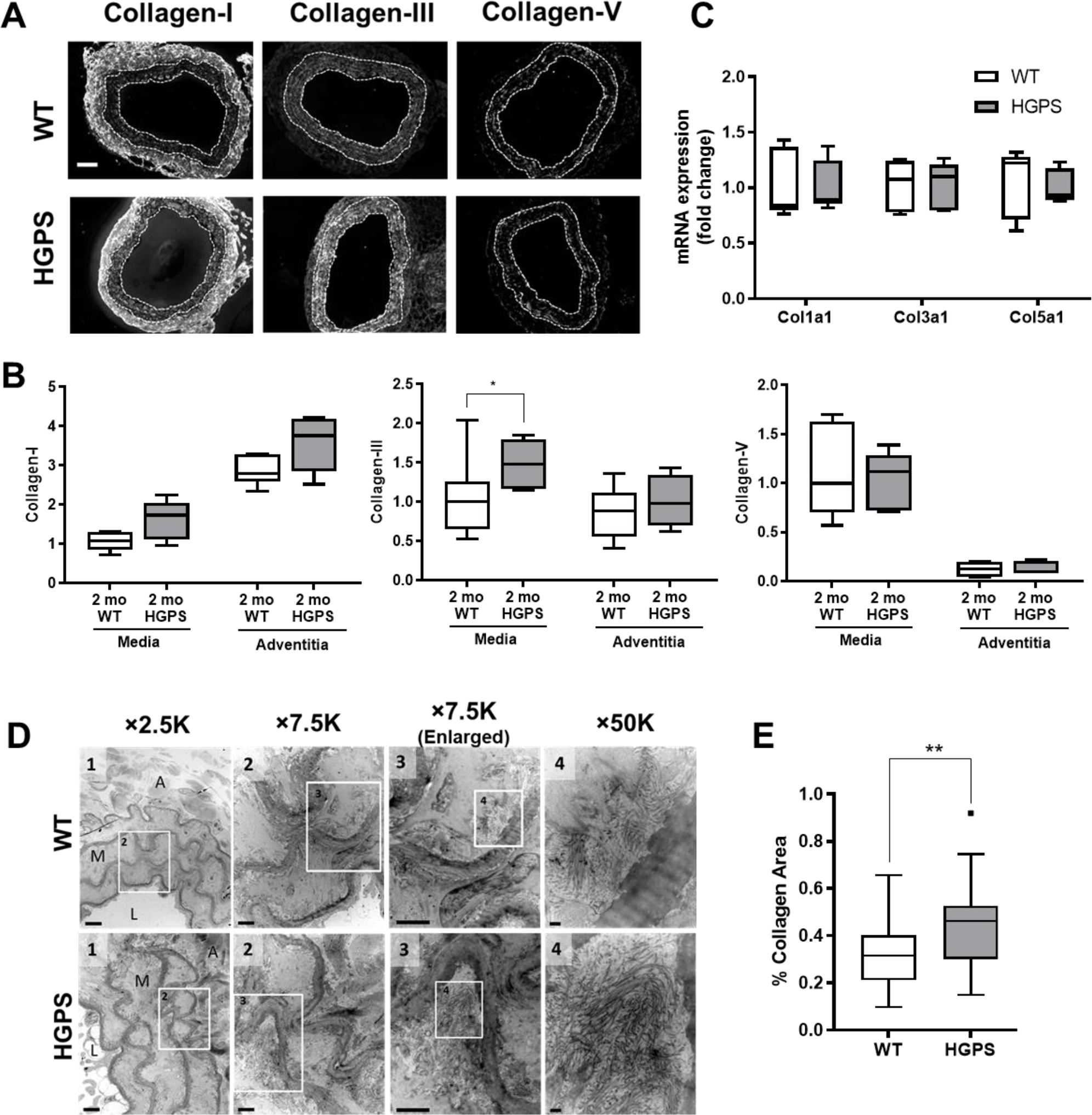
HGPS carotid arteries display alterations in abundance of fibrillar collagens. **(A)** Paraffin-embedded carotid artery cross sections from 2-month WT and HGPS mice (n=5-7 mice per genotype) were immunostained for collagens I, III, and V; scale bar 50 μm. The medial layer is outlined in dotted lines. **(B)** Results were quantified and plotted relative to the median signal intensity of 2-month WT media. Statistical significance was determined by Mann-Whitney tests. **(C)** RNA isolated from aortas of 2-month WT and HGPS mice was analyzed by RT-qPCR (n=5 per genotype). Results were analyzed by Mann-Whitney tests for each mRNA. **(D)** Carotid arteries from 2-month WT and HGPS mice were sectioned and imaged by TEM. Representative images are shown at the indicated magnification. Numbers in each image correspond to the boxed ROI in the preceding image. In image 1, the lumen (L), media (M), and adventitia (A) of the artery are noted (scale bar = 6 μm). Image 2 shows multiple elastin folds, and image 3 shows a 2-fold enlargement of an individual fold used to quantify collagen area as described in Supplemental Methods (scale bar= 2 μm). In image 4, individual collagen fibrils within the elastin fold are readily distinguishable (scale bar = 200 nm). **(E)** Collagen area in elastin folds of WT and HGPS carotid arteries was quantified as described in Methods (n=3 mice, 30 elastin folds). Statistical significance was determined by Mann-Whitney test.

We then used Transmission Electron Microscopy (TEM) to search for potential differences in collagen abundance between WT and HGPS arteries that might escape detection by traditional immunostaining. (Fig. 2D). The high resolution of TEM revealed that collagen in the medial layer of both WT and HGPS mouse carotid arteries was restricted to the elastin folds (Fig. 2D), similar to observations in human aortas (Greenberg 1986). The area of collagen in these folds was slightly increased in HGPS (Fig. 2E).

### Upregulated Lysyl Oxidase dominates early arterial ECM remodeling in HGPS

In contrast to the results with collagens, immunostaining revealed a pronounced increase in expression of LOX in the carotid arteries of HGPS mice as compared to WT; most of the LOX was expressed in the medial layer (Fig. 3A). Comparison of LOX expression with time showed that median LOX expression in the media increased ~7-fold between 1- and 2-months of age in the HGPS mice, but insignificantly in the WT controls (Figs. 3B). In contrast, the abundance of adventitial LOX was similar between both WT and HGPS mice (Fig. 3 C). We then assessed transcript abundance for each of the LOX family members by performing RT-qPCR on extracted medial layers of thoracic aortas from 2-month male WT and HGPS mice. We observed an overall trend of increased abundance of all LOX isoforms, and statistically significant increases in LOX and LOXL4 (Fig. 3D). A comparison of delta-CT values of WT mice aorta indicate that LOX is the predominant isoform in the medial layer (Fig. S6).

**Fig. 3.**
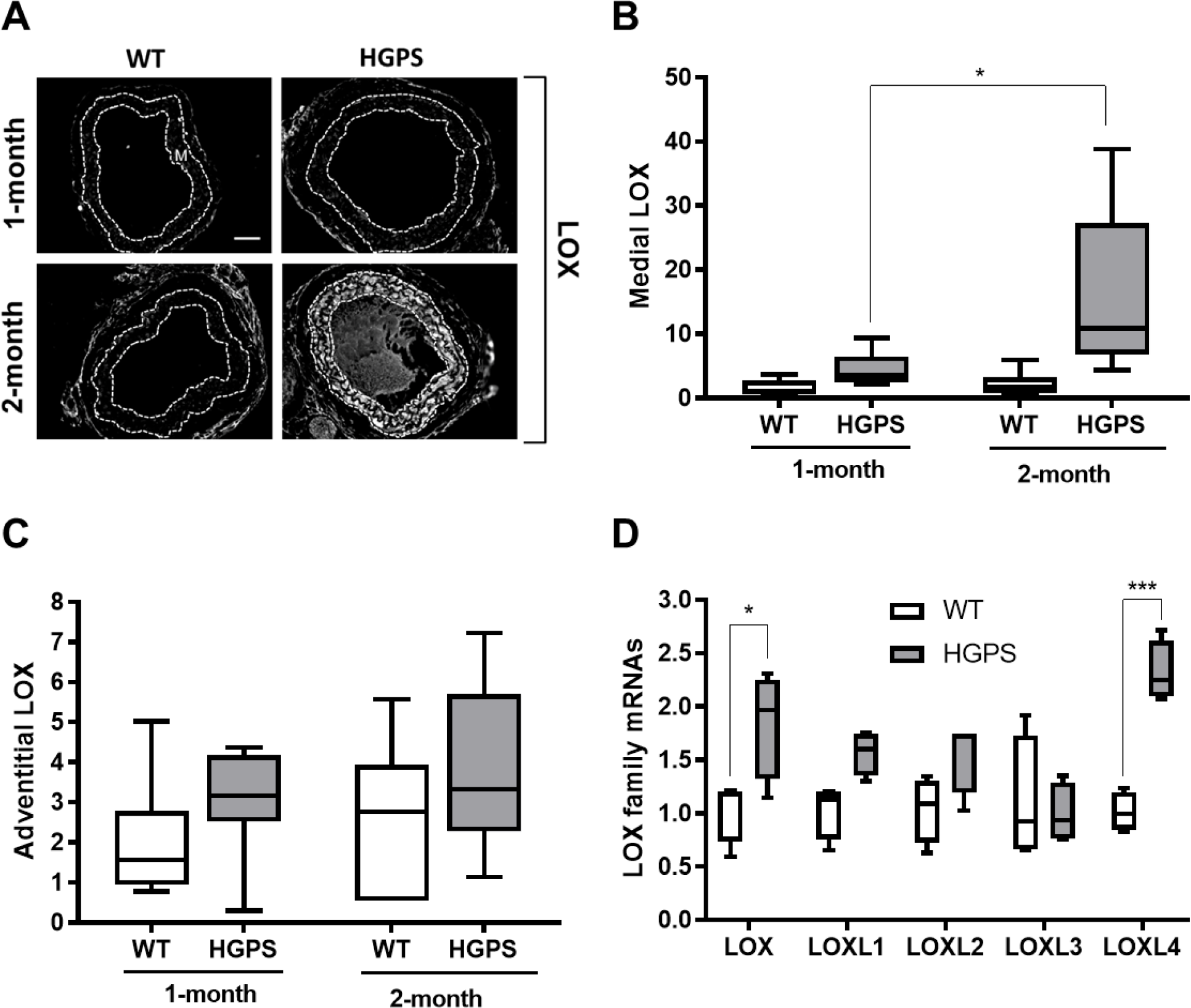
LOX abundance increases between 1- and 2-months in HGPS mice. **(A)** Representative LOX immunostaining of paraffin-embedded carotid artery sections from 1- and 2-month WT and HGPS mice (n=7-9 mice per age and genotype). The medial layer (M) is outlined in dotted lines. Scale bar = 50 μm. **(B-C)** Quantified LOX signal intensities from the medial and adventitial layers of WT and HGPS carotid arteries, respectively; results were normalized to the median signal intensity of the 1-month WT medial layer. Statistical significance between 1- and 2-month old WT mice and 1- and 2-month old HGPS mice was determined by Mann-Whitney test. **(D)** LOX isoform mRNA levels in the medial layer of 2-month WT and HGPS aortas were quantified by RT-qPCR (n=4 experiments, with 2 aortas pooled per experiment). Statistical significance was determined by two-way ANOVA followed by Holm-Sidak post-tests.

### The *Lysyl Oxidase inhibitor BAPN corrects circumferential arterial mechanics in HGPS mice*

Because we observed increased mRNAs for multiple arterial LOX isoforms in HGPS, we treated mice with the pan-LOX inhibitor β-aminopropionitrile (BAPN) (Tang et al. 1983; Jung et al. 2003; Kim et al. 2003; Lee & Kim 2006; Rodriguez et al. 2010) to determine the role of LOX in the early arterial stiffening of HGPS. We administered BAPN between 1-2-months of age, when the levels of medial LOX are increasing (Fig. 3) and used a concentration and injection regime that we previously showed to be effective in altering fibrillar collagen structure and reducing arterial stiffness (Kothapalli et al. 2012). Consistent with our previous work demonstrating that BAPN does not affect arterial collagen abundance (Kothapalli et al. 2012), TEM analysis showed that the area of collagen within elastin folds was similar without or with BAPN treatment (Fig. S7).

Biaxial inflation-extension tests examined the effect of LOX inhibition on arterial mechanics of the HGPS arteries. Administration of BAPN to HGPS mice barely affected axial mechanics, with the BAPN-treated cohort showing a similar IVS to vehicle-treated controls, and only a slightly increased axial stress (Figs. 4A-B). In contrast, BAPN had a major effect on circumferential stiffness and resulted in a circumferential stress-stretch curve that closely resembled the WT controls (Fig. 4C). Consistent with arterial softening, BAPN restored the stress-stretch relationship of HGPS arteries to near WT levels, and this effect was associated with a slight increase in the inner radius (Fig. 4D) and slight decrease in wall thickness (Fig. 4E). Administration of BAPN to WT mice had little effect on arterial mechanics other than a small decrease in axial stress at high stretch (Fig. S8). Collectively, the elevated expression of LOX coincident with premature circumferential arterial stiffening, and the correction of this stiffness defect by BAPN establishes premature LOX induction as a critical mechanism underlying premature arterial stiffening in HPGS. Our analysis further shows that HGPS targets the expression of LOX mRNA as well as the mRNAs for less abundantly expressed LOX family members.

**Fig. 4.**
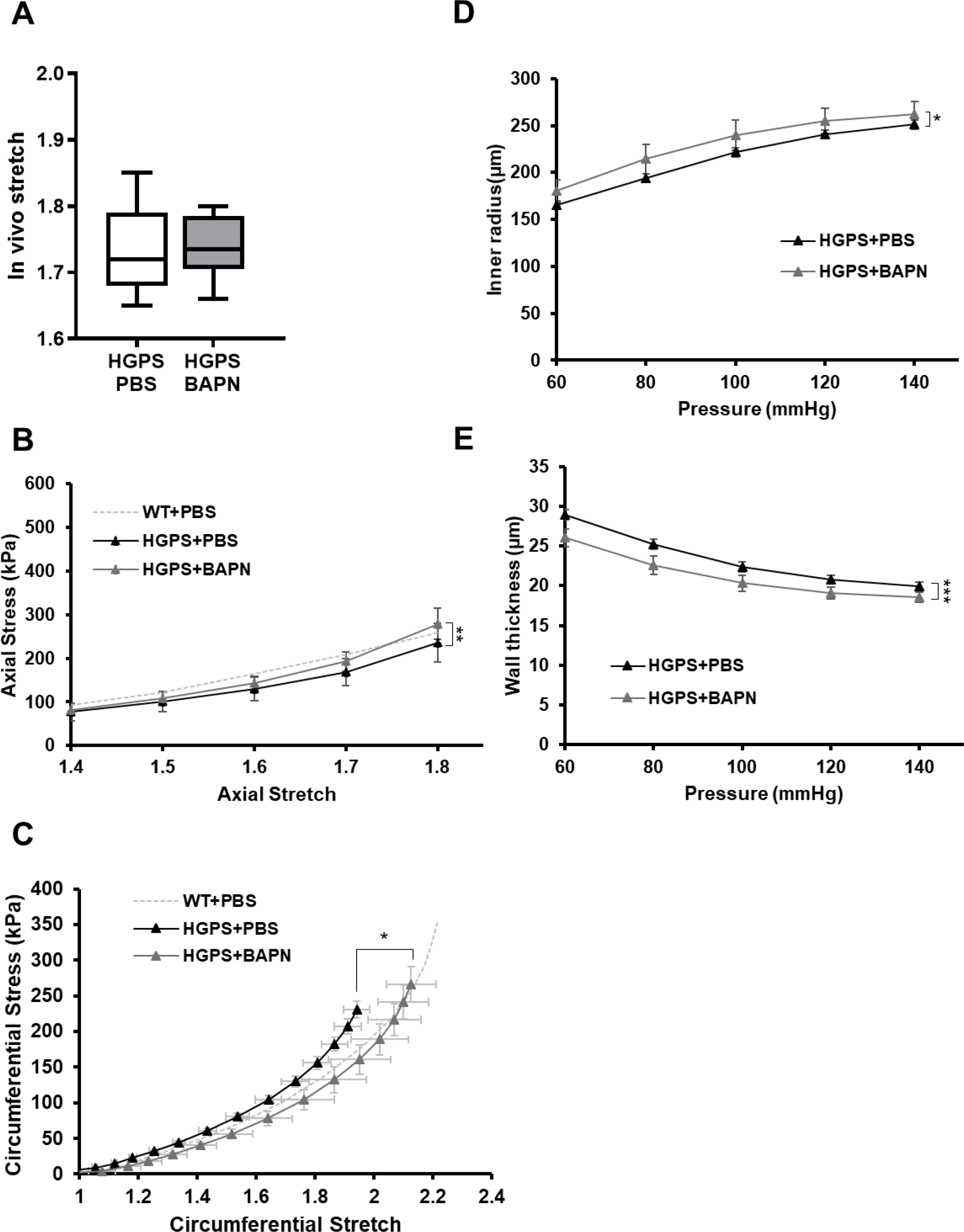
LOX inhibitor BAPN reduces circumferential stiffening of HGPS carotid arteries. HGPS mice aged to 1-month were treated with BAPN or PBS (vehicle) for 30 days followed by pressure myography of isolated carotid arteries from HGPS mice treated with PBS (n=7) or BAPN (n=6). **(A)** *In vivo* stretch of carotid arteries. **(B)** Axial stress-stretch curves of carotid arteries; results show means ± SD. **(C)** Circumferential stress-stretch curves show means ± SE. Statistical significance between PBS-treated and BAPN-treated HGPS mice was determined by two-way ANOVA. Grey dashed lines in panels B and C show the stress-stretch relationship of 2-month WT arteries as reference (see Fig. S8 for primary data). **(D-E)** Measured inner radii and wall thickness, respectively, of carotid arteries from PBS or BAPN-treated HGPS mice; results show means ± SE with statistical significance determined by two-way ANOVA.

### Lysyl Oxidase expression is increased coincident with circumferential arterial stiffening in normal aging

As many parallels have been drawn between HGPS and normal aging, we also evaluated levels of the fibrillar collagens and LOX in aged (24-month) WT mice. The carotid arteries of the aged WT mice and young (2-month) HGPS mice were similar in showing only small changes in medial collagen III and collagen-V (compare Figs. 5A and S9 with Fig. 2A-B). However, aged WT mice showed a statistically significant increase in adventitial collagen-I that was not seen in HGPS (Fig. 5A). A time course study from 2-24 months revealed that adventitial collagen-I begins to increase at ~12-months and continued to increase to 24-months in WT mice (Fig. 5B). The same analysis showed that LOX expression increased with time in both the medial and adventitial layers and reached maximal levels between 12 and 24-months of age (Fig. 5B-C). In fact, this is the age range that shows increased arterial stiffening in WT mice (Brankovic et al. 2019).

**Fig. 5.**
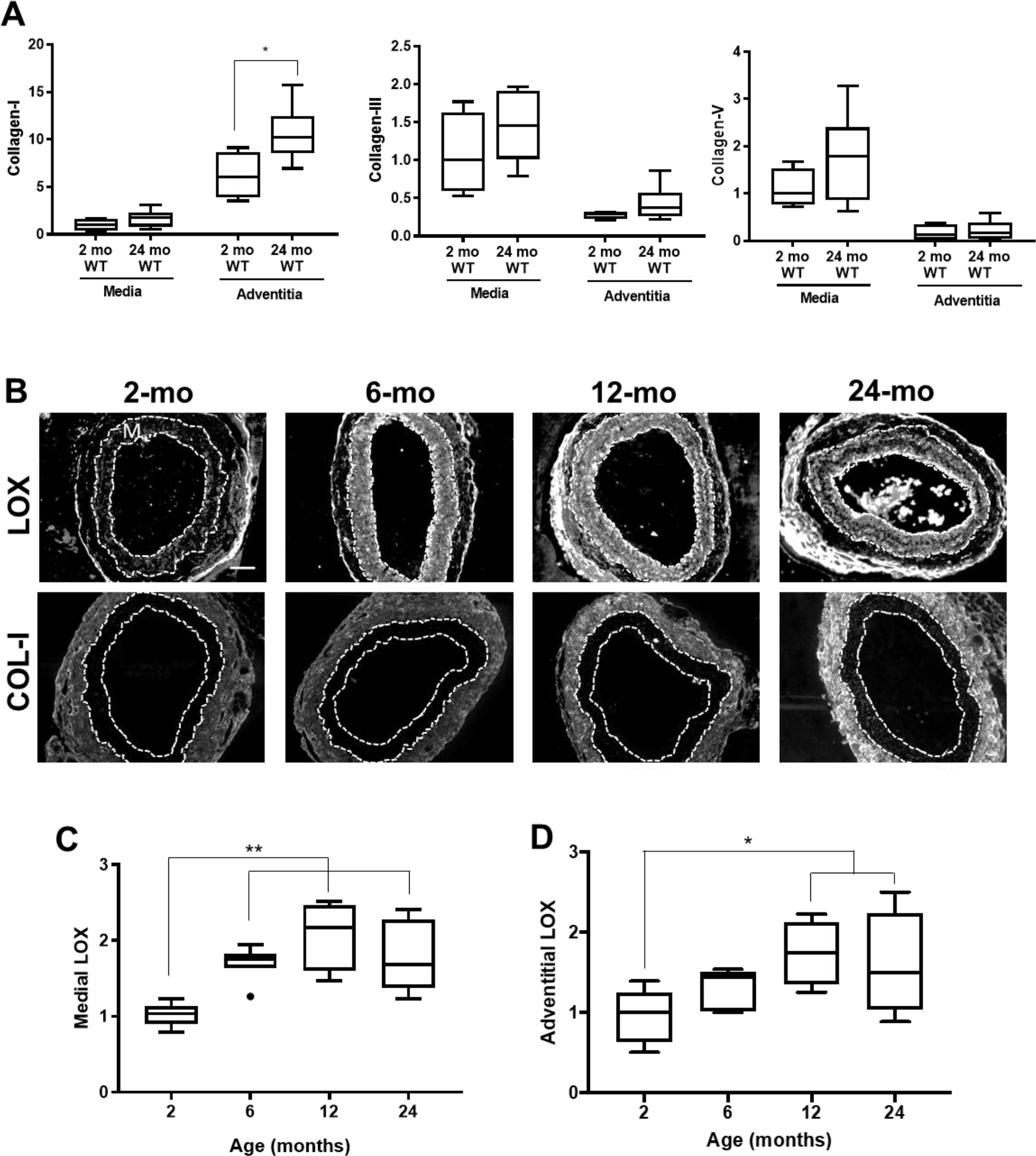
Increased LOX abundance in carotid arteries of aged WT mice. C57BL/6 (WT) mice were aged from 2-24 months. **(A)** Quantification of medial and adventitial collagen-I, III, and V from immunostaining of carotid artery cross sections of 2- and 24-month mice (n=4-5 mice). Results were normalized to the median signal intensity of 2-month WT media. Statistical significance was determined by Mann-Whitney tests. Representative images are shown in Fig. S9. **(B)** Representative images of LOX and collagen-I immunostaining of paraffin-embedded carotid artery cross sections from 2-month (n=8), 6-month (n=7), 12-month (n=4), and 24-month (n=9) WT mice. The medial layer (M) is outlined in dashed lines; scale bar = 50 μm. **(C)** Medial LOX signal intensity was quantified and normalized to the median signal intensity of the 2-month WT medial layer. **(D)** The analysis in panel C was repeated for the adventitial layer. Statistical significance in panels C-D was determined by one-way ANOVA relative to the 2-month old mice and followed by Holm-Sidak post-tests.

## DISCUSSION

We show here that the carotid arteries of HGPS mice stiffen prematurely and that this stiffening is preferentially circumferential. The expression of several arterial collagens is elevated in old HGPS mice (del Campo et al. 2019). However, our results show that the initiation of arterial stiffening in HGPS is much more closely associated with increased medial LOX expression. This increase in LOX protein is associated with increases in the levels of LOX mRNA, an effect that was most pronounced for LOX itself but also seen in the other LOX family members. Importantly, pharmacologic intervention with the pan-LOX inhibitor, BAPN, corrected the aberrant arterial mechanics of HPGS mice, thereby establishing LOX upregulation as an underlying and causal mechanism in the early arterial stiffening of HGPS.

While we did not see pronounced increases in arterial fibrillar collagens early in HGPS, TEM revealed a slight increase in collagen area within the medial elastin folds of HGPS arteries. As adventitial collagen abundance was similar in WT and HGPS mice, the medial collagen in these folds may be the target of the elevated medial LOX in HGPS. Indeed, previous studies have reported that the media layer contributes more toward circumferential stiffening while the adventitial layer contributes more to axial mechanics (Kohn et al. 2015). Since the expression of collagen genes can increase with ECM stiffness (Kothapalli et al. 2012), the LOX-mediated increase in arterial stiffness described in our studies here with very young HGPS mice may lead to an eventual induction of other SMC collagens as seen when HGPS mice are aged (del Campo et al. 2019).

As progerin has been detected in aged tissues, cells, and atherosclerotic lesions (Scaffidi & Misteli 2006; Olive et al. 2010; McClintock et al. 2007), there has been significant interest in the idea that progerin-like splicing may contribute to phenotypes of natural aging as well as drive early aging in HGPS. Indeed, both 2-month HGPS and 24-month WT carotid arteries display increased expression of medial LOX and increased circumferential stiffness. However, the axial mechanics of 2-month male HGPS carotid arteries are mostly intact, while 24-month male WT carotids display clear axial stiffening. This distinction implies that, in addition to the common effect on LOX abundance, there must also be inherent differences between the arterial stiffening process in normal aging versus premature aging in HGPS.

Others have recently reported increased LOX and LOXL2 in old (20-22 month) versus young (3-4 month) aortas of WT mice (Steppan et al. 2019). This work complements our findings of increased LOX expression in HGPS carotid arteries, and the joint findings indicate that increased LOX activity with age is likely to be a general feature of large artery stiffening. In our work, LOX, rather than LOXL2, is the major arterial LOX family member. Our spatial analysis also shows that LOX is induced in both the arterial medial and adventitial layers with natural aging whereas LOX induction is largely restricted to the media in HGPS. Thus, while overall LOX induction is a common feature of arterial stiffening in both natural aging and HGPS, its distinct spatial expression patterns also distinguish natural aging and HGPS.

Others have reported that administration of sodium nitrate to HGPS mice (del Campo et al. 2019) and administration of Lonafarnib, a farnesyltransferase inhibitor, to HGPS children reduce arterial stiffness (Gordon et al. 2012; Gordon et al. 2018). Furthermore, the reduction in arterial stiffness by lonafarnib was associated with an increased lifespan of HGPS children (Gordon et al. 2018). These studies did not distinguish between effects on axial versus circumferential mechanics, nor are the mechanisms by which nitrate and lonafarnib reduce arterial stiffness fully clear. Nevertheless, these studies and this report collectively indicate that decreasing arterial stiffness could lessen the burden of cardiovascular disease in HGPS. Our results further indicate that targeting circumferential arterial mechanics, potentially though LOX inhibition, may be an important consideration when developing mechanically inspired therapeutics for HGPS.

## EXPERIMENTAL PROCEDURES

### Mice and artery isolation

WT C57BL/6 mice were purchased from Jackson Labs and aged to 24-months. LMNA^G609G/+^ mice were generously provided Dr. Carlos Lopez-Otin (Universidad de Oviedo, Oviedo, Spain). Mice were fed a chow diet *ad libitum*. At the appropriate age, the mice were sacrificed by CO_2_ asphyxiation, the left carotid artery was immediately removed, stripped of most fat, and used for pressure myography as outlined below. The remaining arteries were perfused in situ with PBS. The right carotid artery was then removed, cleaned in PBS and then fixed in either Prefer for paraffin-embedding or TEM fixative (see below). The descending aorta was isolated from the end of the aortic arch to the diaphragm, cleaned as above, and used for the preparation of RNA (see Supplemental Methods).

### Biaxial extension-inflation tests using a pressure myograph

Arterial mechanics were determined on a DMT 114P pressure myography with force transducer largely as described (Brankovic et al. 2019). Freshly isolated carotid arteries from WT and HGPS mice (Table S1) were secured to 380 μm (outer diameter) cannulas using silk sutures; blood was cleared, and any remaining fat was removed. Once mounted, the arteries were visualized by light microscopy, and the unloaded/unpressurized arterial wall thickness and inner radius was measured at the axial length where the artery transitioned from being bent to straight (Table S1). Arteries were brought to a stretch of 1.7 and pressurized to 100 mm Hg for 15 min with medical air (Airgas). The arteries were then preconditioned by cyclic pressurization three times from 0–140 mmHg in 1-min increments. Unloaded (unstretched and unpressurized) vessel wall thickness and outer diameter were measured in multiple sections after preconditioning and averaged for post-test data analysis.

In vivo stretch (IVS) was determined using force-length tests as described (Brankovic et al. 2019; Ferruzzi et al. 2013). Briefly, the carotid arteries were axially stretched in 10% increments at three constant pressures (90, 120, and 140 mm Hg). Equilibrium force was recorded for each stretch and pressure, and the intersection of the three force-stretch curves was defined as the IVS. Circumferential stiffness, loaded inner radius, and loaded wall thickness were determined from pressure-outer diameter tests with samples at their IVS and pressurized in 10 mm Hg, 30-sec steps from 0–140 mm Hg before returning the artery to 0 mm Hg (Brankovic et al. 2019). This test was performed three times, and the three stress-stretch curves were averaged. We confirmed the validity of our IVS determinations by measuring axial force through the circumferential tests, and we excluded samples where axial force varied from the mean by >25% with pressure. See Supplemental Methods for methods of data analysis.

### Transmission Electron Microscopy (TEM) analysis of collagen structure

Carotid arteries for TEM were fixed overnight at 4°C in 0.1M sodium cacodylate buffer, pH7.4 containing 2.5% glutaraldehyde and 2.0% paraformaldehyde. After subsequent buffer washes, the samples were post-fixed in 2.0% osmium tetroxide for 1 hour at room temperature and rinsed in water prior to *en bloc* staining with 2% uranyl acetate. Briefly, after dehydration through a graded ethanol series, the tissue was infiltrated and embedded in EMbed-812 (Electron Microscopy Sciences, Fort Washington, PA). Thin sections were stained with uranyl acetate and lead citrate and examined with a JEOL 1010 electron microscope fitted with a Hamamatsu digital camera and AMT Advantage image capture software. Images of artery sections were taken at increasing magnification; collagen abundance was evaluated as described in Supplemental Methods.

### In vivo treatment with BAPN

Mixed sex WT and HGPS mice (~1 month of age) were injected peritoneally with BAPN (Sigma A3134; 333 mg/kg) or an equal volume of vehicle (control). BAPN was dissolved in PBS and injected in a volume of 0.2 ml per day until the mice reached 2-months of age. No dramatic alterations in mouse behavior, appearance, or weight loss was observed during the injection period. At 2-months of age, the mice were sacrificed, carotid arteries were isolated, and unloaded wall thickness and inner radii were determined (see Table S2). The samples were then analyzed by pressure myography as described above.

### Statistical Analysis

All statistical analysis was performed using Prism software (Graphpad). For stress-stretch curves, stretches were either set by the experimenter (axial tests) or calculated using Equation 3 (circumferential). Differences in axial stress were analyzed by two-way ANOVA relative to 2-month WT carotid arteries.Two-way ANOVAs were also used to analyze differences in circumferential stretch and stress relative to 2-month WT arteries; the least significant parameter is shown in the figures. Significance of changes in inner radius and wall thickness were also determined through two-way ANOVA. Two-tailed Mann-Whitney tests were used to evaluate significance of *in vivo* stretch, TEM, RT-qPCR, and immunofluorescent images of arterial sections unless stated otherwise. Statistical significance for all graphs is demarcated by *(p<0.05), **(p<0.1), ***(p<0.001). Axial stress-stretch curves show means ± SD. Circumferential stress-stretch curves as well as inner radius and wall thickness results were determined from triplicate determinations per sample and are therefore presented as means ± SE. Results are represented either as line graphs with means and either SD or SEM as described in the legends or as box and whisker plots with Tukey whiskers.

## ACKNOWLEDGEMENTS

We thank the Electron Microscopy Research Laboratory at the University of Pennsylvania for preparing arterial tissue for TEM. This work was supported by NIH grant AG047373, the Progeria Research Foundation and the Center for MechanoBiology, a National Science Foundation Science and Technology Center under grant agreement CMMI15457. RvK was supported by NIH grants T32-GM008076 and F31-HL142160

## Author Contributions

Experiments were designed by RvK and RKA. Experiments were performed by RvK, SB, IR, EAH, KB, and PC. Data analysis and statistical testing was conducted by RvK, SB, IR, KB, and RKA. Animal management and drug administration was conducted by RvK and EAH. The manuscript and figures were prepared by RvK and RKA.

