## Supplemental Data for "Premature arterial stiffening in Hutchinson-Gilford Progeria Syndrome linked to early induction of Lysyl Oxidase (LOX) and corrected by LOX inhibition"

**Supplemental Methods**

*Carotid artery immunostaining and analysis.* Paraffin-embedded sections (5 µm) of freshly isolated carotid arteries from 2-month WT and 2-month LMNA$^{G609G/G609G}$ were hydrated before antigen unmasking (Vector Labs, H3300). Sections were washed in PBS three times before blocking with 2% BSA in PBS for 15 min, incubated overnight at 4ºC with antibodies directed to p16$^{INK4A}$ (Proteintech, 10883-1-AP; 1:50 dilution), collagen-I (Southern Biotech, 1310-01; 1:400 dilution), collagen-III (Proteintech, 22734-1-AP; 1:400 dilution), collagen-V (Abcam, ab7046; 1:250 dilution), and LOX (Santa-Cruz, sc32409 selective for the LOX isoform; 1:50 dilution). Replicate sections were incubated in parallel with isotype-matched control antibodies. All samples were washed three time with PBS before incubation with a 1:100 dilution of Alexa 594-conjugated isotype-matched secondary antibody (Lifetech donkey anti-goat A11058 or Invitrogen Goat anti-rabbit A11012) for two hours at room temperature. Sections were then washed three times in PBS followed by addition of Dapi (1:1000 dilution in PBS). Slides were briefly washed in PBS and then water before mounting with SlowFade Gold (Thermo, S36936). Results were visualized with a Nikon Eclipse 80i microscope with a QI-Click Qimaging camera. Carotid arteries were imaged at 20x magnification.

Images were quantified using ImageJ. The media of each section was traced using the polygon drawing tool, and its raw integrated intensity was divided by the area of the outlined media to obtain relative fluorescence intensity. The procedure was then repeated for the adventitial layer. Relative fluorescence intensity values were then plotted relative to the median fluorescence intensity value of the 2-month WT. Median fluorescent intensity values were passed through a Grubb's test. Background intensity, as determined from the isotype-matched control antibodies, was negligible (see Fig S5). Results are presented as box plots with Tukey whiskers.

*Histological analysis of arterial sections.* Arteries were fixed after excision in Prefer (Anatech #414), embedded in paraffin and 5-µm sections were stained. Staining with Hematoxylin (Fisher,

SH30-500D) and Eosin (Fisher, SE22-500D) used standard procedures. Apoptosis was determined by immunostaining for cleaved caspase-3 (Cell Signaling Technologies SignalStain Apoptosis IHC Detection kit, #12692S) according to manufactures instructions with tumor xenograft sections as positive controls. To determine calcium deposition, sections were deparaffinized and hydrated followed by addition of a 5% solution of Alizarin Red S (Sigma, A5533) pH 4.2 for 30 minutes with calcified bone tissue as positive control. Arterial elastin layers were visualized by autofluorescence using a cyan filter on a Nikon Eclipse 80i Fluorescence microscope.

*RT-qPCR*. Descending aortas from 2-month WT and HGPS mice were isolated and either stored immediately in RNAlater (Qiagen) for analysis of collagen mRNAs or stripped of adventitia for analysis of LOX mRNAs. To strip adventitia from the aorta, a cleaned isolated aorta was incubated in 1 mg/ml Type 2 collagenase (Worthington Biochem, LS004174) in Hanks Balanced Salt Solution (Lifetech, 14170-112) for 10 minutes at 37$^{o}$C. The adventitial layer was carefully peeled off before storing the aorta in RNAlater. Total RNA was isolated with the RNeasy Fibrous Tissue Mini Kit (Qiagen) according to manufacturer's instructions and using 0.3 ml buffer RLT per aorta. Reverse transcription reactions contained 200-500ng of total RNA. Ten to fifteen percent of the cDNA was subjected to qPCR with the following primer-probe sets from Applied Biosystems: Col1a1 (Mm00801666_g1), Col3a1 (Mm00802300_m1), Col5a1 (Mm00489299_m1); LOX (Mm00495386_m1), LOXL1 (Mm01145738_m1), LOXL2 (Mm00804740), LOXL3 (Mm01184865_m1), LOXL4 (Mm00446385). The primer-probe set for 18s rRNA has been described in Klein et al., 2007 (Klein et al. 2007). Results were normalized to 18S rRNA, and changes in mRNA abundance were calculated using the ddCT method.

*Myograph Data analysis.* Measurements of intraluminal pressure, force, and outer diameter were converted into stress-stretch curves using equations 1–4 where *l* and *L*=loaded and unloaded vessel lengths, respectively (μm), $a_i$ and $A_i$=loaded and unloaded inner radii, respectively (μm), $a_o$ and $A_o$=loaded and unloaded outer radii, respectively (μm), $h$ and $H$ = loaded and unloaded vessel wall thickness, respectively (μm), $P$= intraluminal pressure (mm Hg), and $f_T$=axial force (nN). Vessel wall thickness was calculated in the post-test analysis as described (Brankovic et al. 2019) with the standard assumption that the sample was incompressible.

Equation 1: Axial stretch $(\lambda_z)=\frac{l}{L}$

Equation 2: Axial stress $(\sigma_z)= \frac{Pa^2\pi + f_T}{\pi h(2a + h)}$

Equation 3: Circumferential stretch $(\lambda_\vartheta)=\frac{a + h/2}{A + H/2}$

Equation 4: Circumferential stress $(\sigma_\vartheta)=\frac{Pa i}{h}$

*Quantification of collagen within elastin folds.* TEM images of WT and HGPS carotid arteries were taken at 7500x magnification to visualize the elastin folds (Fig S10; A1). The folds were defined as areas where the elastin invaginates, creating a roughly parabolic shape (Fig. S10; A2). Using the polygon Selection Tool in ImageJ, an area was defined by continuously tracing along the two sides of the parabolic shape; that region was then enclosed by connecting the two apexes with a straight line (Fig. S10; A3). This area was added to the ROI Manager to calculate total area of the elastin fold. To calculate the area within the fold containing collagen, the Paintbrush Tool was used to manually black-out areas containing collagen fibers (Fig. S10; A4). The painted area was isolated with the Threshold Tool (Fig. S10; A5) and added to the ROI manager (Fig. S10; A6). The ratio of collagen/total areas was defined as percent collagen in the elastin fold.

**Supplemental Tables**

| | WT (2 mo) male (n=7) | HGPS (2 mo) male (n=6) | WT (24 mo) male (n=5) | WT (2 mo) female (n=7) | HGPS (2 mo) female (n=6) | WT (24 mo) female (n=5) |
|---|---|---|---|---|---|---|
| **Mouse age (days)** | 68 ± 4.9 | 61 ± 1.3 | 721 ± 6.9** | 64 ±1.2 | 63 ± 0.60 | 731 ± 3.7** |
| **Artery unloaded outer diameter (µm)** | 368 ± 7.7 | 332 ± 12* | 412 ± 11* | 340 ± 9.2 | 322 ± 9.3 | 404 ± 11** |
| **Artery unloaded wall thickness (µm)** | 65 ± 1.9 | 61 ± 2.6 | 73 ± 1.2* | 65 ± 1.2 | 66 ± 1.2 | 70 ± 0.27** |

**Table S1. Characteristics of murine carotid arteries.** Values from 2-month and 24-month WT mice and 2-month HGPS mice are shown as means ± SE. A Mann-Whitney test was performed between the 2-month WT and HGPS groups and the 2-month WT and 24-month WT groups within each sex.

| | WT (PBS; n=6) | WT (BAPN; n=5) | HGPS (PBS; n=7) | HGPS (BAPN; n=6) |
|---|---|---|---|---|
| **Mouse age (days)** | 59 ± 2.4 | 61 ± 0.80 | 60 ± 1.3 | 59 ± 1.6 |
| **Artery unloaded outer diameter (μm)** | 367 ± 27 | 342 ± 11 | 331 ± 4.7 | 320 ± 4.9 |
| **Artery unloaded wall thickness (μm)** | 69 ± 0.81 | 69 ± 0.18 | 68 ± 0.33 | 69 ± 0.30 |

**Table S2. Arterial characteristics after BAPN treatment.** Mouse age, unloaded outer diameter, and unloaded wall thickness of the carotid arteries used for pressure myography testing after injections with vehicle (PBS) or BAPN. Results show means ± SE.

**Supplemental Figures**

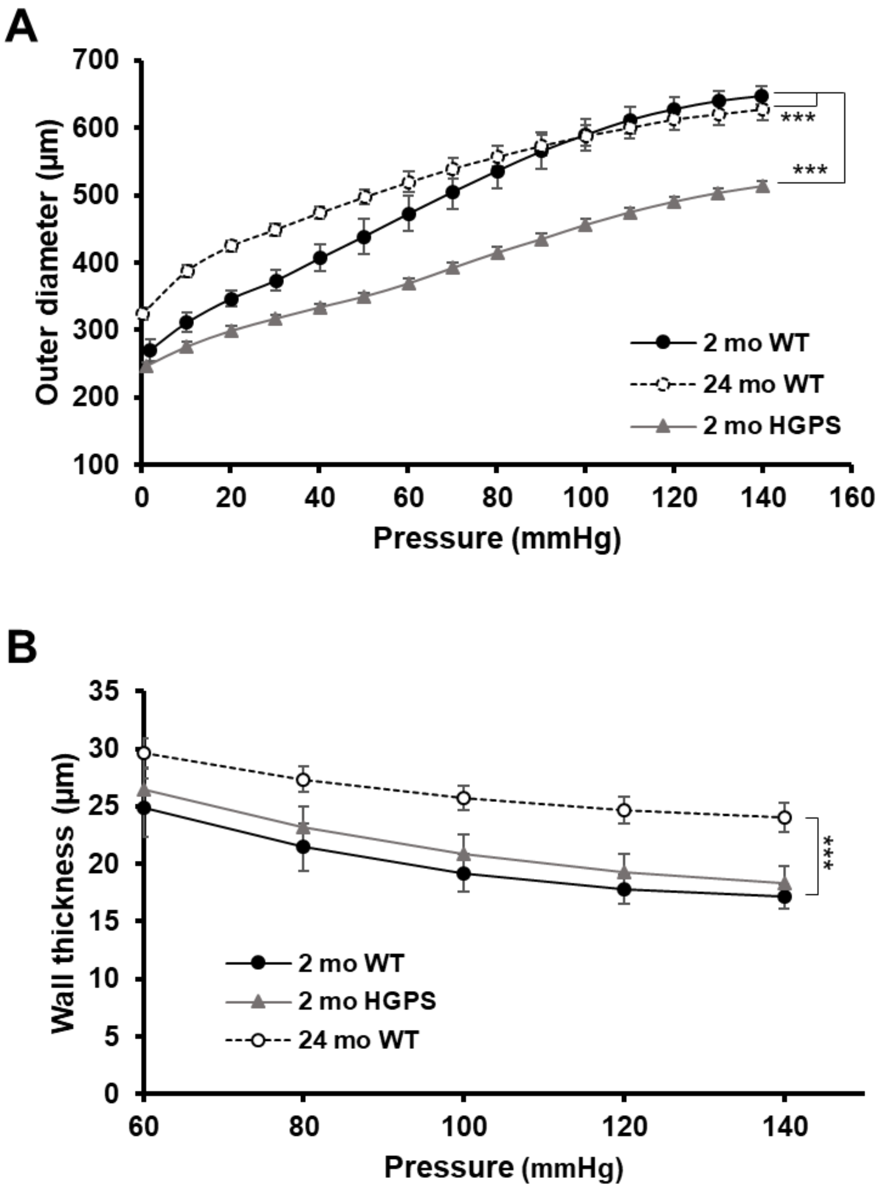


**Figure S1. Changing carotid artery geometry with age and HGPS. (A)** Pressure-outer diameter relationships in 2-month (n=6) and 24-month (n=5) WT mice and 2-month HGPS mice (n=6) as determined by pressure myography. **(B)** Mean carotid wall thickness ± SE. Statistical significance was determined by two-way ANOVA and relative to 2-mo WT mice.

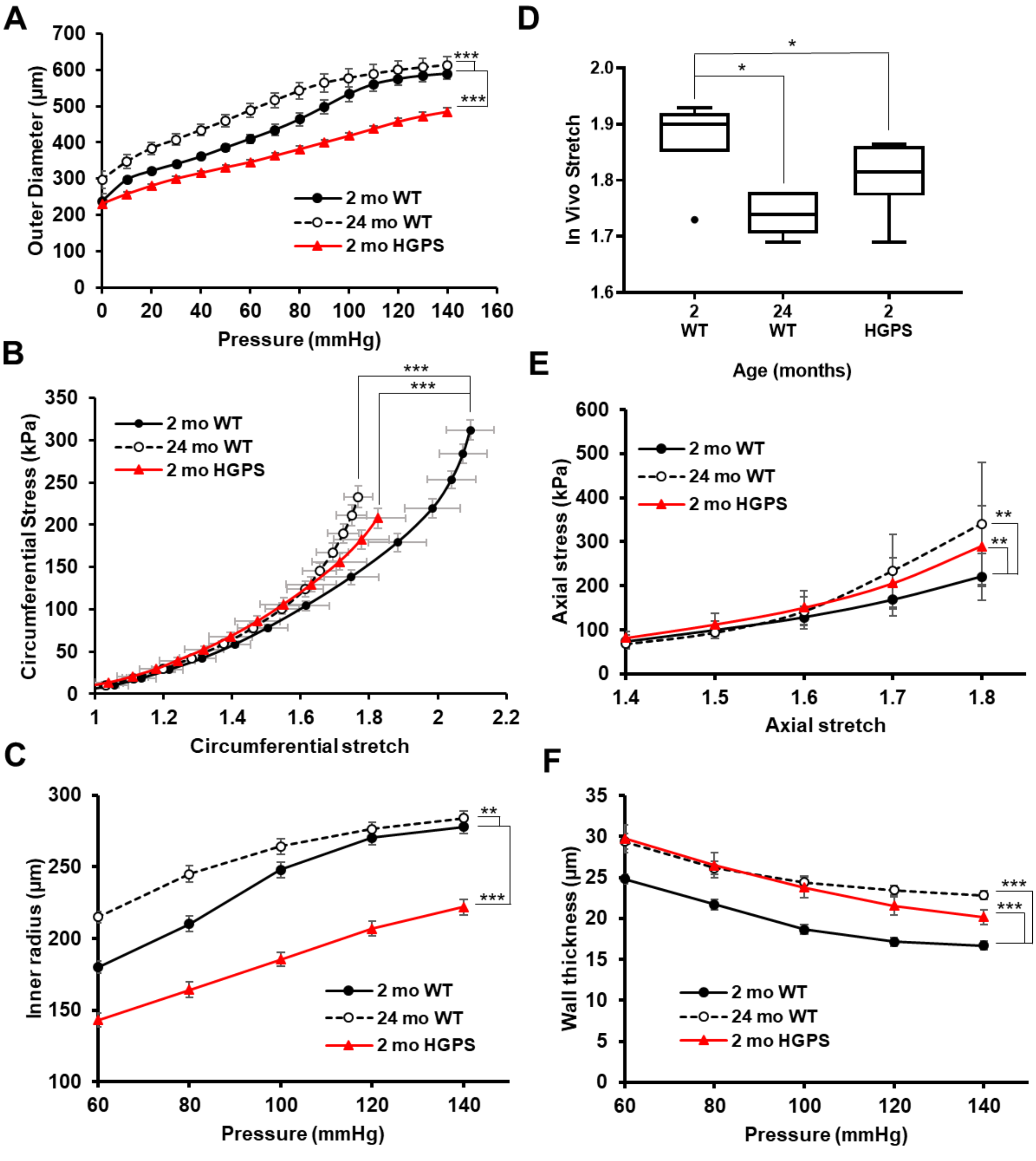


**Figure S2. Mechanical properties of female WT and HGPS carotid arteries.** Carotid arteries from female 2-month WT (n=7), 2-month HGPS (n=6), and 24-month WT (n=5) mice were analyzed by pressure myography. **(A)** Pressure-outer diameter curves. **(B)** Circumferential stress-stretch curves. Results show mean ± SE **(C)** Graph of inner radii displaying mean ± SE. **(D)** Calculated IVS. **(E)** Axial stress-stretch curves performed at 80-90 mm Hg with mean ± SD. **(F)** Graph of wall thickness displaying mean ± SE. Statistical significance was determined either by Mann-Whitney test (panel D) or by two-way ANOVA (panels A-C,E-F) relative to 2-month WT arteries.

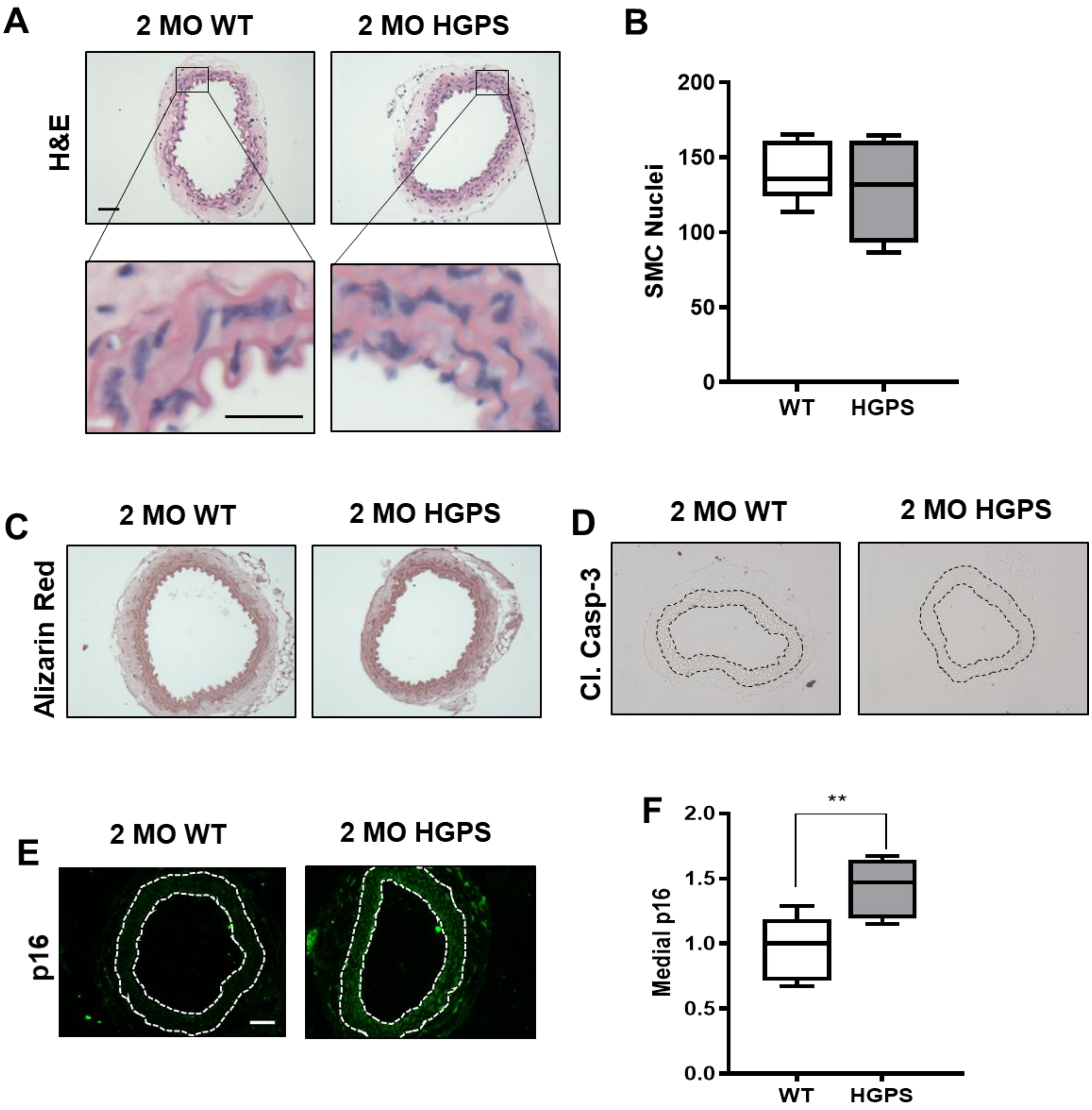


**Figure S3. Histology of carotid arteries from 2-month WT and HGPS mice. (A)** Carotid artery cross sections were stained with H&E; scale bar = 50 μm and inset bar = 25 μm. **(B)** The number of medial SMC nuclei was quantified from H&E images (n=4 per genotype with 4 sections analyzed per mouse). **(C)** Cross sections were stained with Alizarin Red (n=3-5 per genotype), **(D)** immunostained for cleaved caspase-3 (n=3-5 per genotype), or **(E)** immunostained for the senescence marker, p16INK4A. **(F)** Quantification of medial p16INK4A normalized to the median signal intensity of carotid sections from 2-month WT mice; (n=6-10 per genotype). Statistical significance was determined by Mann-Whitney test.

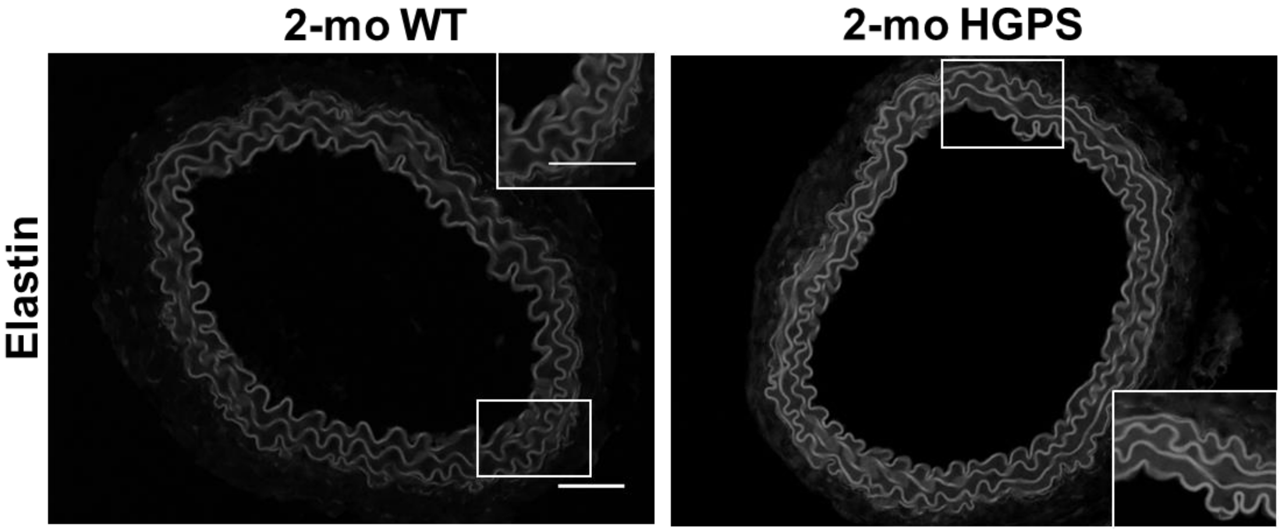


**Figure S4. Elastic laminae of WT and HGPS carotid arteries.** Representative elastin autofluorescence images of 2-month WT and HGPS carotid artery cross sections (n=7-9 per genotype). Scale bar = 50 μm.

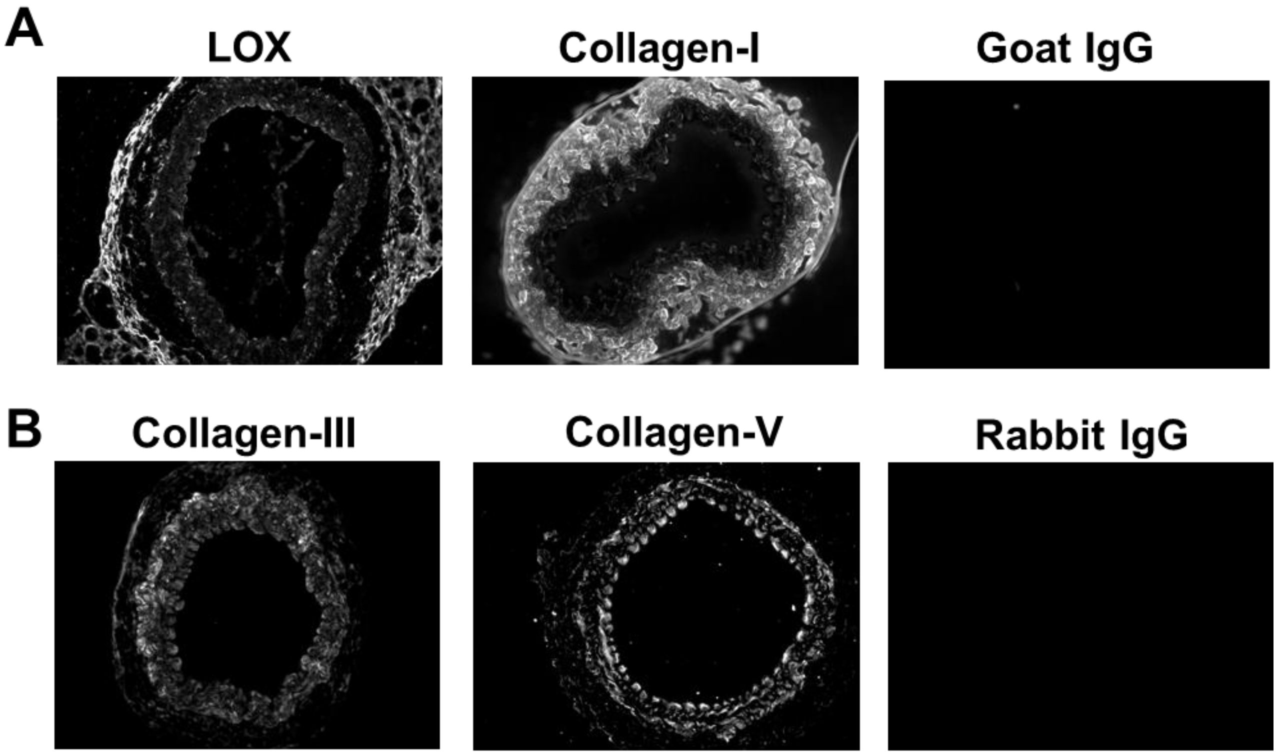


**Figure S5. Controls for immunostaining.** Representative images of background signals for carotid artery cross sections immunostained with targeted and isotype-matched control goat (LOX and COL1) and rabbit (COL3 and COL5) antibodies.

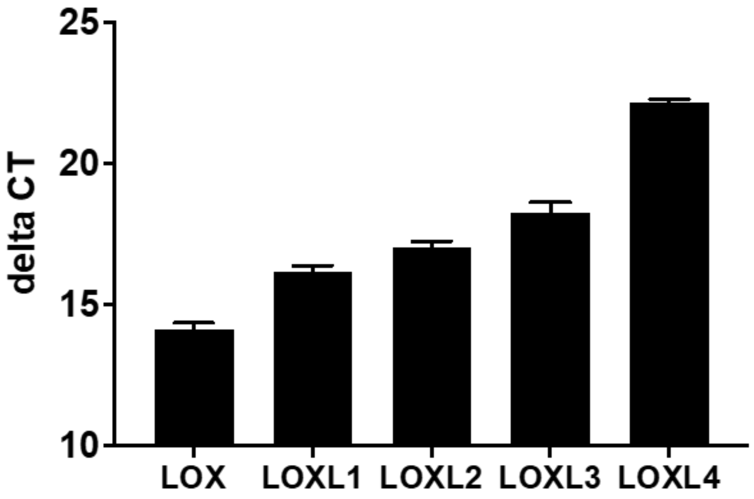


**Figure S6. Relative mRNA levels of LOX family members in WT aortas.** For each sample, two 2-month WT aortas were stripped of adventitia (see Supplemental Methods) combined and used for isolation of total RNA. The relative abundance of the LOX family member was determined by RT-qPCR. Results show delta-CT values plotted as mean ± SE (n=4).

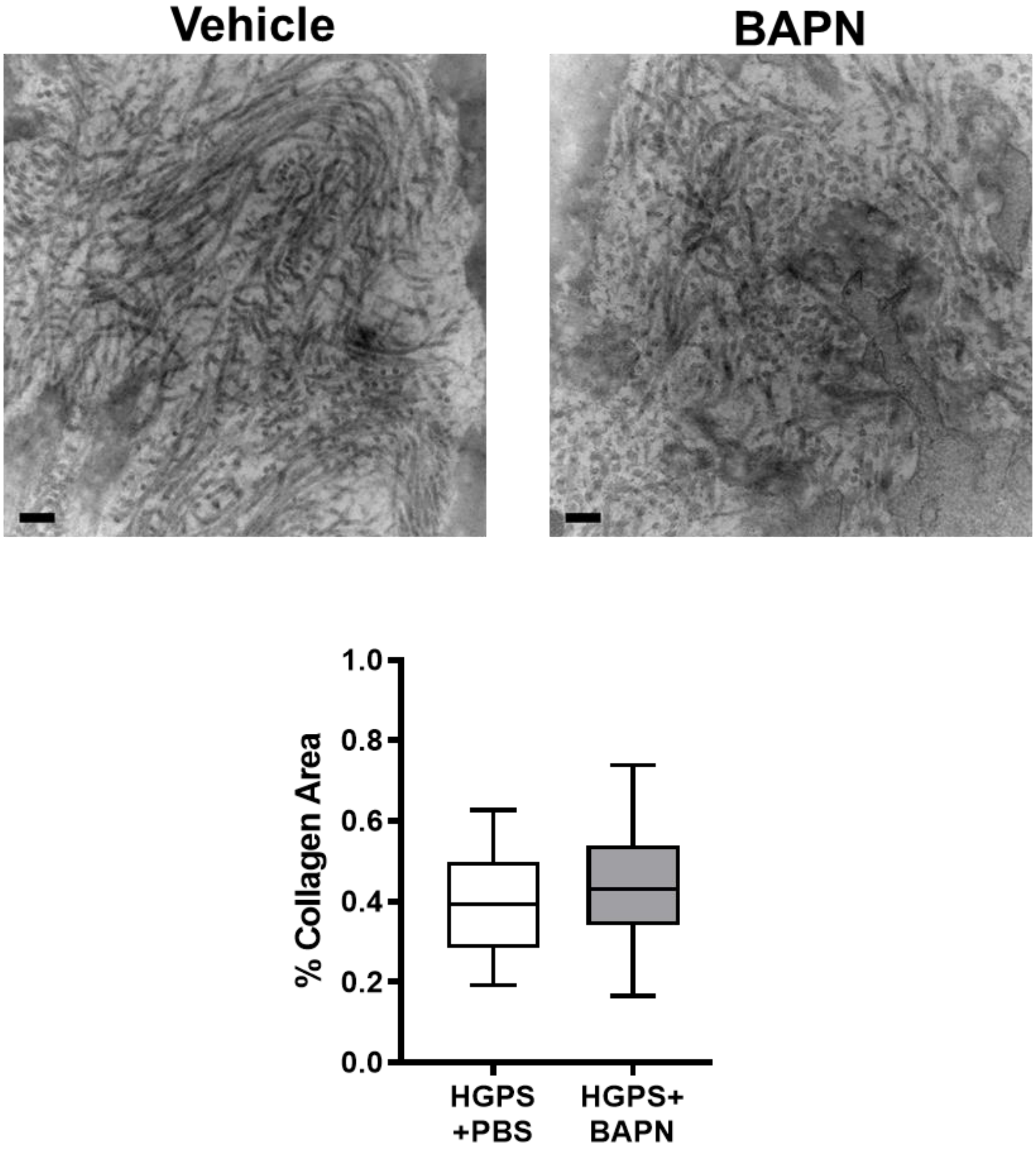


**Figure S7. TEM images of elastin folds of HGPS carotid arteries. (A)** Representative TEM images (50,000x magnification) of cross sections from 2-month HGPS carotid arteries that had been treated with vehicle (PBS) or BAPN as described in Methods. Scale bar = 200 nm. **(B)** Quantification of collagen area within the elastin folds as described in Supplemental Methods and Fig. S10.

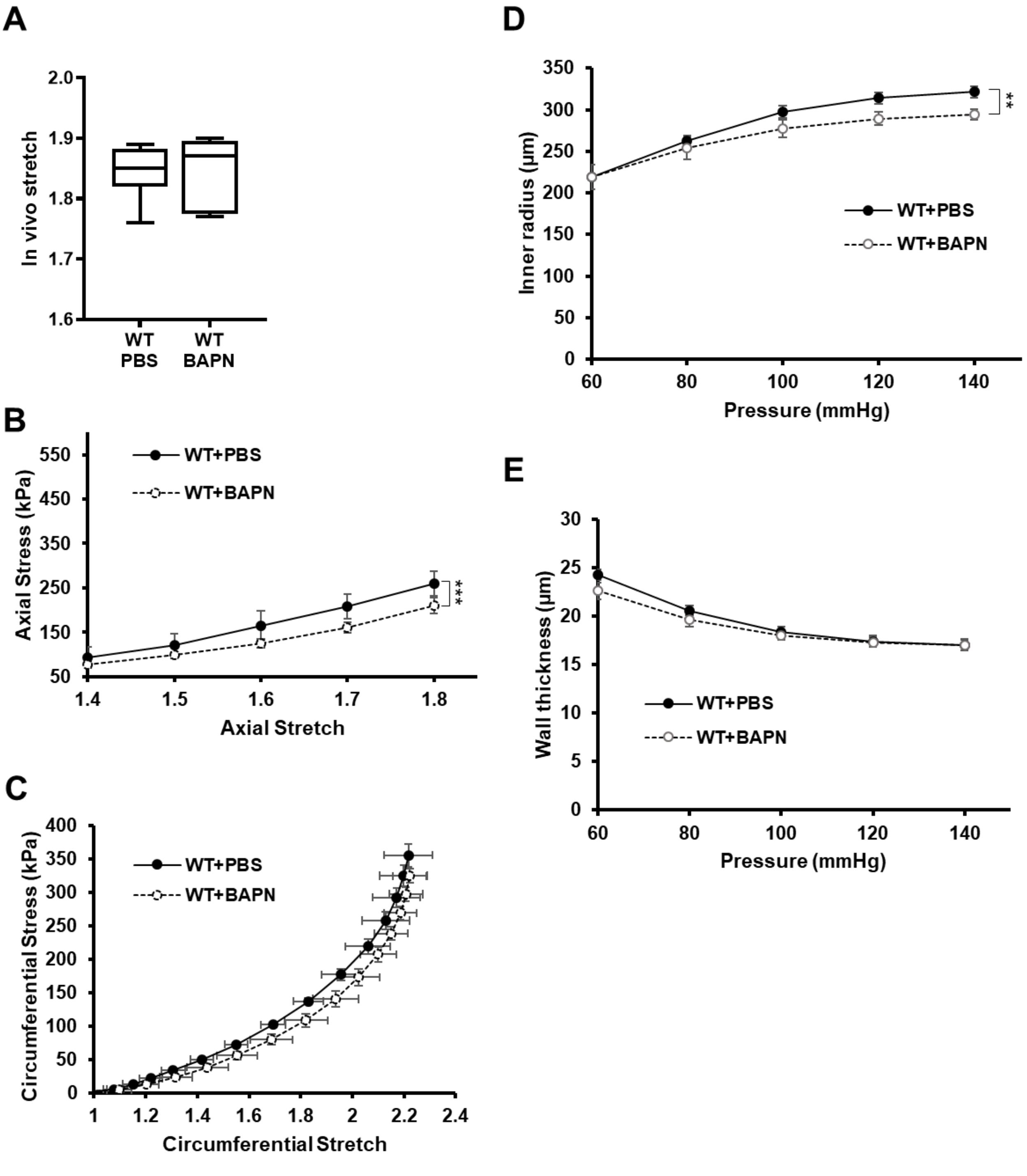


**Figure S8. Mechanical parameters of PBS and BAPN-treated mice. (A)** IVS of PBS- and BAPN-treated WT (n=6 and 5, respectively) mice. **(B-C)** Axial and circumferential stress-stretch curves, respectively, for PBS- and BAPN-treated WT mice. **(D)** Inner radii and **(E)** wall thickness were measured with changing pressure. Results in B show means ± SD. Results in C-E show means ± SE. Statistical significance in B-E was determined by two-way ANOVA.

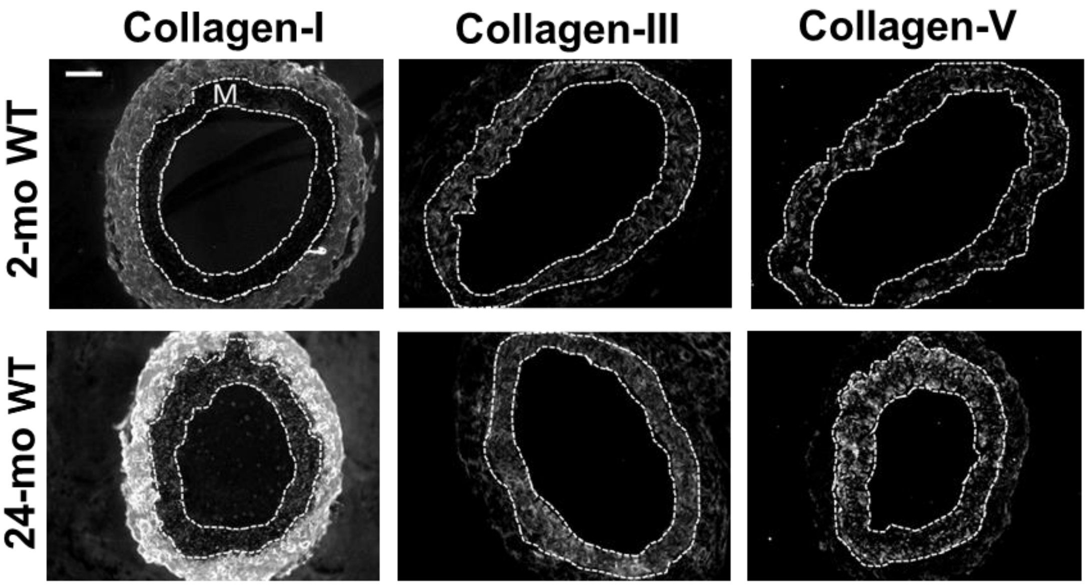


**Figure S9.** Representative images of paraffin-embedded carotid artery cross sections from 2-month and 24-month WT (n=4-7 mice per age group) immunostained for collagens I, III, or V. The media layer (M) is outlined in dotted lines. Scale bar = 50 μm.

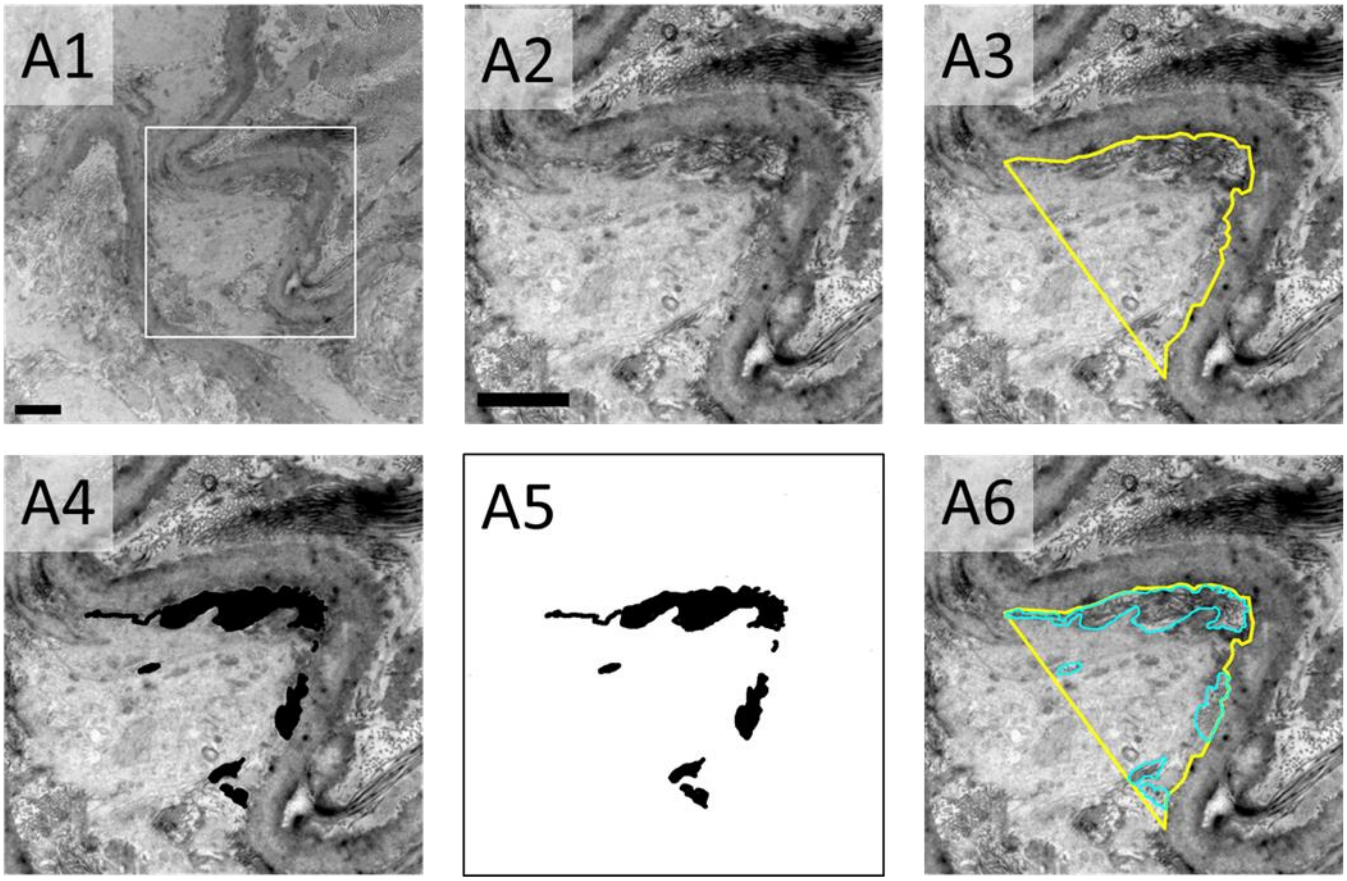


**Fig S10. Quantification of collagen abundance in elastin folds.** Representative images from a mouse carotid artery cross section processed in ImageJ to quantify the area of collagen within an elastin fold. The boxed region in image A1 is shown in images A2-A6. Scale bar = 2 µm. See Supplemental Methods for details.
